# Fluctuations of fMRI activation patterns reveal theta-band dynamics of visual object priming

**DOI:** 10.1101/148635

**Authors:** Bingbing Guo, Zhengang Lu, Jessica E. Goold, Huan Luo, Ming Meng

## Abstract

The brain dynamically creates predictions about upcoming stimuli to guide perception efficiently. Recent behavioral results suggest theta-band oscillations contribute to this prediction process, however litter is known about the underlying neural mechanism. Here, we combine fMRI and a time-resolved psychophysical paradigm to access fine temporal-scale profiles of the fluctuations of brain activation patterns corresponding to visual object priming. Specifically, multi-voxel activity patterns in the fusiform face area (FFA) and the parahippocampal place area (PPA) show temporal fluctuations at a theta-band (~5 Hz) rhythm. Importantly, the theta-band power in the FFA negatively correlates with reaction time, further indicating the critical role of the observed cortical theta oscillations. Moreover, alpha-band (~10 Hz) shows a dissociated spatial distribution, mainly linked to the occipital cortex. These findings, to our knowledge, are the first fMRI study that indicates temporal fluctuations of multi-voxel activity patterns and that demonstrates theta and alpha rhythms in relevant brain areas.

## INTRODUCTION

To efficiently interact with an ever-changing environment, it has been proposed that the brain generates perceptual predictions about forthcoming stimuli based on previously primed hypothesis (Heekeren, Marrett, and Ungerleider 2008, Bar 2009, Gorlin et al. 2012, Rao and Ballard 1999, Engel, Fries, and Singer 2001, Summerfield et al. 2006). In circumstances containing multiple predictions, it would make sense that multiple predictions are dynamically coded and organized to maximize the efficiency of anticipation. Consistent with this notion, recent human behavioral results, using a visual priming paradigm, reveal that competing perceptual predictions are temporally coordinated in competing theta-band waves (i.e., being conveyed in various phase of the theta-band oscillations) (Huang, Chen, and Luo 2015). However, little is known about the neural mechanisms underlying this rhythmic coordination.

Theta-band (3-8 Hz) neuronal oscillations have been demonstrated in perception, memory, and attention (Lisman and Idiart 1995, VanRullen and Koch 2003, Busch and VanRullen 2010, Landau and Fries 2012, Luo et al. 2013, Fiebelkorn, Saalmann, and Kastner 2013, Landau et al. 2015). Here we focus on the neural basis of the rhythmic activities in visual object priming, by examining the temporal dynamics in several related brain areas. First, since a face or a house image was used to be the prime/probe stimulus, we would expect to find theta-band rhythms in the fusiform face area (FFA (Kanwisher, McDermott, and Chun 1997)) and the parahippocampal place area (PPA (Epstein and Kanwisher 1998)). Alternatively, theta rhythms might be also found in early sensory processing brain areas if the theta oscillations are due to rate limitations on sensory sampling. Finally, it is also possible that the rhythm might reflect rate constraints of attentional selection and thus would be revealed in high-level brain areas such as parietal and frontal cortex.

Neural oscillations are ubiquitous (Buzsáki 2006) and have been widely studied with electroencephalography (EEG) and magnetoencephalography (MEG) (Luo et al. 2013, Landau et al. 2015) in human subjects. Here, we employ a novel method that uses functional magnetic resonance imaging (fMRI) to investigate neural dynamics with millimeter-level spatial resolution across the whole human brain. Specifically, we combined fMRI, multi-voxel pattern decoding (Haxby 2012), and a time-resolved behavioral priming paradigm (Huang, Chen, and Luo 2015, Landau and Fries 2012, Fiebelkorn, Saalmann, and Kastner 2013, Song et al. 2014) to assess the fine spatio-temporal profile. In an object priming experiment, a masked prime (i.e., a face or a house image) initially activates a corresponding perceptual prediction, which is then compared to a subsequent probe (i.e., a face or a house) that is either congruent or incongruent with the perceptual prediction triggered by the prime (Huang, Chen, and Luo 2015). Critically, we varied trial-by-trial stimulus onset asynchrony (SOA) between prime mask and probe in small steps of 20 ms, from 200 ms to 780 ms. Thus, time-resolved profiles of the dependent variables (i.e. behavioral measurements and fMRI responses) can be reconstructed as a function of mask-to-probe SOA in steps of 20 ms (corresponding to a 50 Hz sampling frequency), representing the fine temporal course of the prediction conveying processes triggered by the prime. Moreover, by examining the temporal relationships between congruent and incongruent conditions, we could also study the multi-prediction coordination process. Recent studies used the time-resolved behavioral measurement in combination with transcranial magnetic stimulation (TMS) and MEG (Dugue, Roberts, and Carrasco 2016, Wutz et al. 2016). We were the first to take this approach to fMRI with multi-voxel pattern decoding.

Figure 1A shows the experimental design. In each trial, a 150-ms probe was preceded by a 33-ms priming stimulus, which was backward masked by a 100-ms mask stimulus. Participants were asked to maintain fixation on a cross displayed in the center of the screen and to make speeded responses to a probe stimulus (detecting a face or house). The prime and probe were either congruent (prime is a face, probe is a face; prime is a house, probe is a house) or incongruent (prime is a face and probe is a house; or vice versa). For each participant, there were 12 repetitions for each of the four prime-probe conditions at each of the 30 SOAs (from 200 to 780 ms in steps of 20 ms). To avoid potential low-level effects of retinotopic adaptation, the size of the prime and probe were different. We also presented face stimuli and house stimuli at different image contrasts to facilitate the decoding of experimental conditions in brain regions of interest (ROIs), including Brodmann area 17 (BA17). Moreover, participants were instructed to use their left and right hands, respectively, to make house and face responses. Therefore, fMRI activity in motor cortex was also tied to corresponding experimental conditions for the majority of correct response trials.

**Figure 1.**
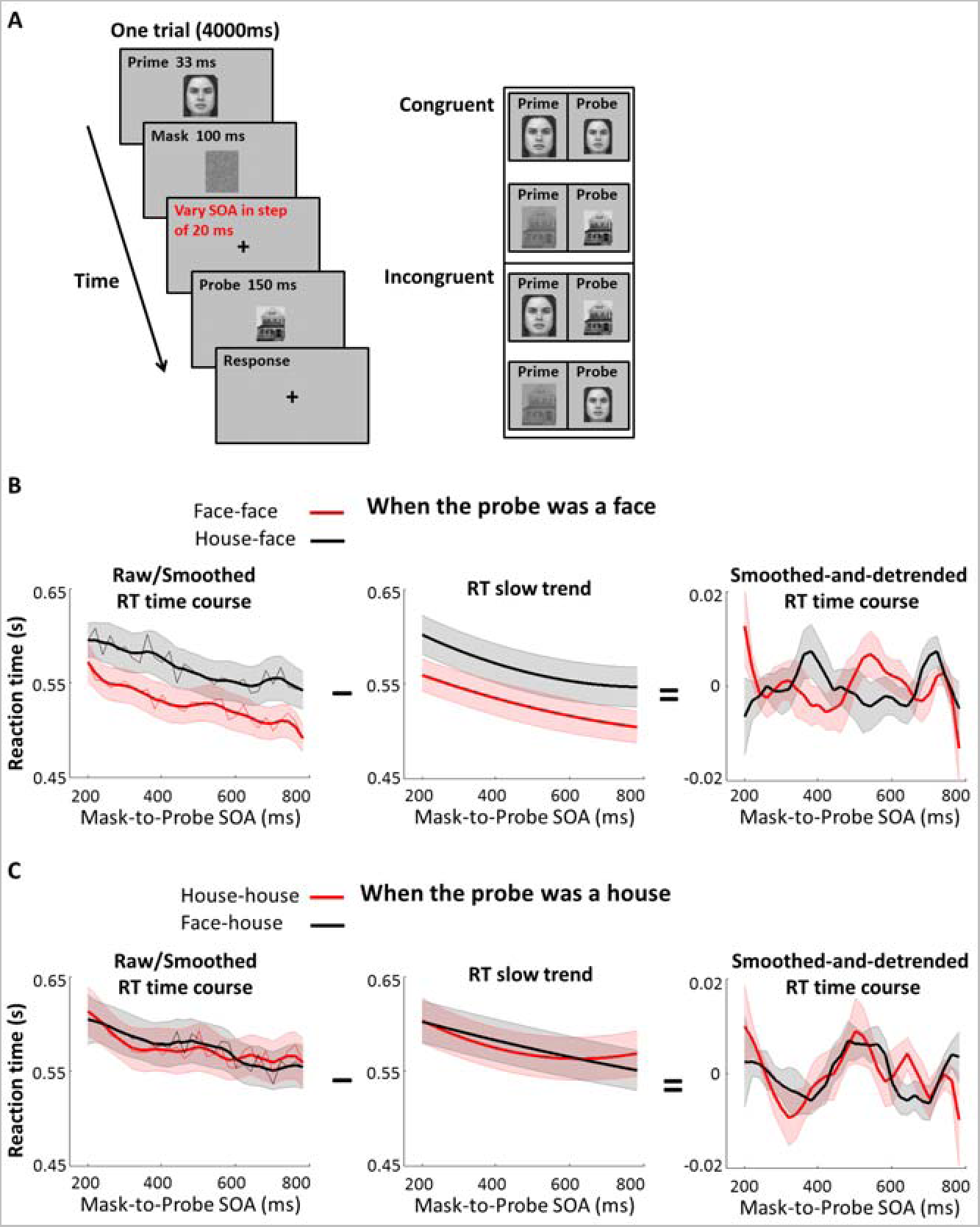
Experiment design and behavioral results. (A) Experiment design. **Left:** For each trial, a 150-ms probe was preceded by a 33-ms priming stimulus, which was backward masked by a 100-ms mask stimulus. Critically, the mask-to-probe stimulus onset asynchronies (SOAs) ranged from 200 ms to 780 ms in steps of 20 ms. **Right:** The stimuli included two images, a high-contrast face and a low-contrast house. Prime and probe stimuli were either the same (congruent) or different (incongruent), except that the probe was always smaller than the prime. (B) Behavioral results of when the probe was a face. **Left:** Average raw RT time courses as a function of mask-to-probe SOA (200-780 ms in steps of 20 ms) for congruent (red, thin line) and incongruent (black, thin line) conditions. Average smoothed (60 ms bin) RT time courses (*n* =18, mean ± SEM) as a function of mask-to-probe SOA, for congruent (red, thick line) and incongruent (black, thick line) conditions, which clearly show an overall priming effect. **Middle:** Average RT slow trends across all the participants. **Right:** Average smoothed-and-detrended RT time courses extracted by subtracting slow trends shown in ***Middle*** from smoothed (60 ms bin) RT time courses shown in ***left*** (thick lines). (C) Behavioral results of when the probe was a house.

## RESULTS

### Behavioral results

All participants (n=18) performed well on the task (percent correct: 97±0.54%). For each participant, we excluded reaction times (RTs) that were >3 SD from the mean across all trials. RT time courses were then plotted as a function of mask-to-probe SOAs. For trials in which the probe was a face (Figure 1B), effects of priming were clearly observed in the raw RT time courses averaged across all participants, as the congruent condition (red, thin line) reliably evoked faster RTs than the incongruent condition (black, thin line). To estimate the RT time courses, we calculated smoothed RT time courses for each participants, starting with probes that were presented from 200 ms to 260 ms (60 ms bin) after the mask, then forward this procedure throughout the mask-to-probe SOAs (200 ms to 780 ms) in step of 20 ms. Figure 1B shows the smoothed RT time courses averaged across all participants for congruent condition (red, thick line) and incongruent condition (black, thick line). To better examine the oscillatory pattern, we extracted slow trends of RT time courses in each participant by using 2nd order polynomial fit to raw RT time courses. After removing slow trends from the smoothed RT time courses in each participant, oscillations of the RT time courses are evident (right panel). This observation replicates the previous findings (Huang, Chen, and Luo 2015), suggesting the effectiveness of our approach to study the dynamics of predictive coding in visual object priming.

Due to perhaps individual differences in exact oscillatory frequency, previous behavioral study (Huang, Chen, and Luo 2015) showed the oscillatory effect reducing with increasing SOA in detrended RT time courses. However, no such effect was found in the present study. Note that while oscillation peaks were the largest during 0-200 ms in that study (Huang, Chen, and Luo 2015), the present study focused on SOAs from 200 ms to 780 ms. Although the largest peaks might only occur at the beginning 200 ms of the averaged data, the effect of oscillation apparently lasts longer than that. For trials in which the probe was a house (Figure 1C), no reliable classical priming effect was observed, presumably due to using a low-contrast house image (therefore, much more difficult to detect than a high-contrast face image) and participants may have detected the house probe simply based on it not being a high-contrast face. Accordingly, due to the weak priming effects in house condition, further fMRI analyses focus on trials when the probe was a face. Nonetheless, it is unlikely that our findings are idiosyncratic to face processing, since significant theta-band oscillations were observed for house-probe conditions when we further tested additional subjects with equal contrast face and house images, complementing similar behavioral oscillation results that had been reported for discriminations of other object categories (Drewes et al. 2015).

### Theta-band oscillations in fMRI patterns

fMRI response patterns were analyzed by using multi-voxel pattern analysis (MVPA) for ROIs including areas in the inferior temporal cortex that are implicated for face processing (FFA (Kanwisher, McDermott, and Chun 1997)) and for house processing (PPA (Epstein and Kanwisher 1998)). Activation patterns in the left motor cortex (lMC), right motor cortex (rMC), BA17, and anterior cingulate cortex (ACC) were also analyzed for comparison. For every trial, the probe was decoded as a face or a house by using MVPA based on fMRI activation patterns in these ROIs. MVPA classification accuracies as a function of SOAs are shown in Figure 2.

**Figure 2.**
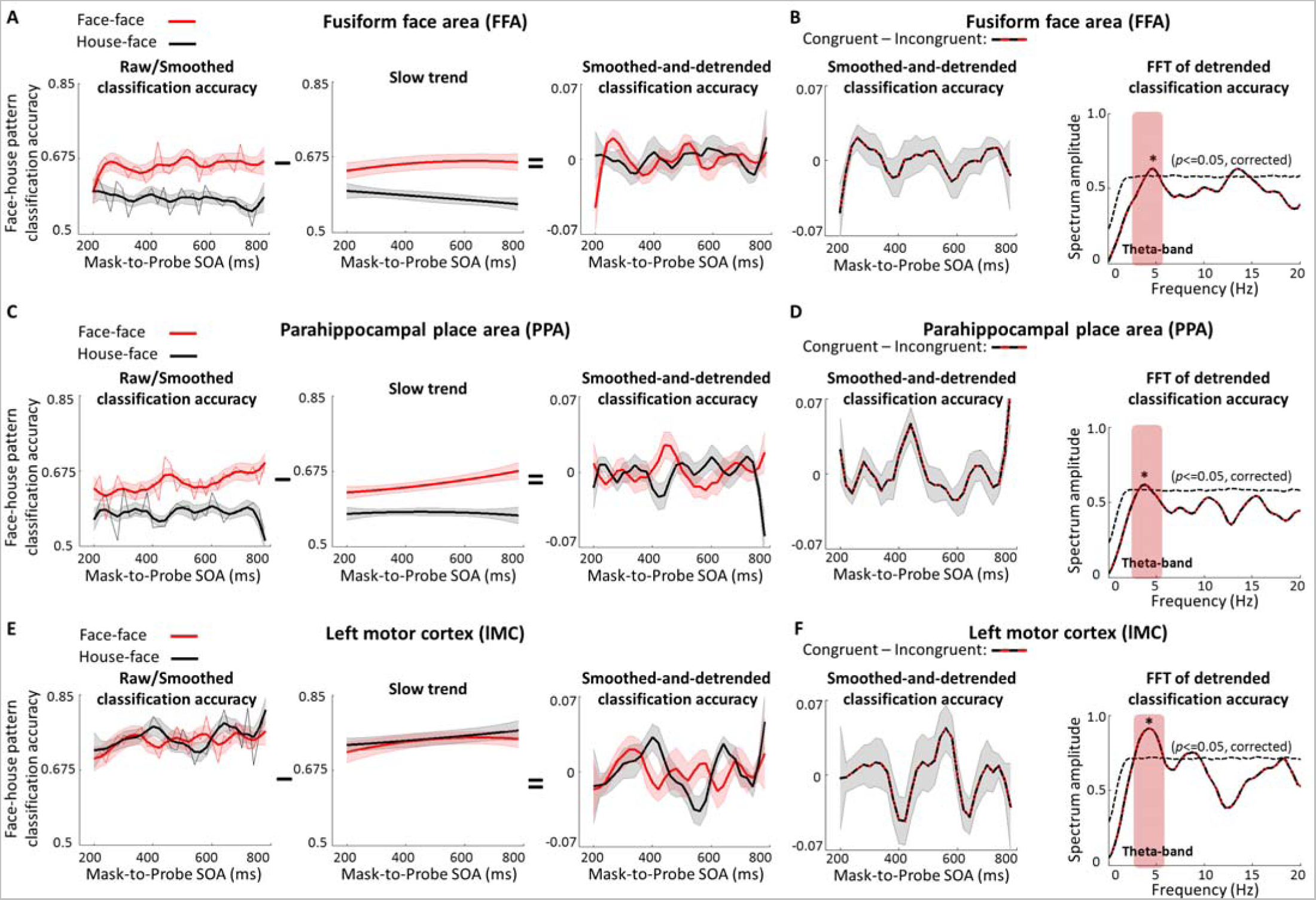
Significant theta-band oscillations of fMRI patterns in the FFA, PPA and lMC. (A) Results in the FFA. **Left:** Average raw classification accuracies as a function of mask-to-probe SOA (200-780 ms) for congruent (red, thin line) and incongruent (black, thin line) conditions. Average smoothed (60 ms bin) classification accuracies (*n*=18, mean ± SEM) as a function of mask-to-probe SOA, for congruent (red, thick line) and incongruent (black, thick line) conditions, which show an overall priming effect. **Middle:** Average slow trends across all the participants. **Right:** Average smoothed-and-detrended classification accuracies extracted by subtracting slow trends shown in ***Middle*** from smoothed (60 ms bin) classification accuracies shown in ***left*** (thick lines). (B) Results of the priming effect (congruent − incongruent) in the FFA. **Left:** Average smoothed-and-detrended classification results of the priming effect. **Right:** Average spectrum for detrended classification accuracies (extracted by subtracting slow trends from the raw classification accuracies) as a function of frequency from 0 to 20 Hz for the priming effect. The statistical threshold of significance (*p* < 0.05, multiple comparison corrected) calculated by performing a permutation test was shown with dashed line. (C) Results in the PPA. (D) Results of the priming effect in the PPA. (E) Results in the lMC. (F) Results of the priming effect in the lMC.

For the raw classification accuracies averaged across all participants, classical priming effects were clearly observed, as the congruent condition (red, thin line) reliably evoked higher classification accuracies than the incongruent condition (black, thin line). To better examine the oscillatory pattern embedded in the MVPA classification accuracies, follow what we have described above for analyzing the behavioral data, we extracted slow trends of classification accuracies. Strikingly, a theta-band rhythm can be clearly seen in the smoothed-and-detrended MVPA results in the FFA, PPA and lMC (Figure 2A, 2C and 2E, right), consistent with the behavioral findings. Furthermore, the oscillations under the congruent and incongruent conditions were in a type of temporally switching relationships (Figure 1B). To examine the out-of-phase relationship between the two conditions, we subtracted the temporal profiles of incongruent conditions (black) from that of congruent conditions (red) for each participant. Oscillations in the effects of priming (congruent − incongruent) averaged across participants are again evident in the FFA, PPA and lMC (Figure 2B, 2D and 2F, left).

To further evaluate the periodicity, the detrended (not the smoothed-and-detrended) priming effects were then converted into frequency domain (after zero padding and application of a Hanning window) (Huang, Chen, and Luo 2015, Song et al. 2014) by using Fast Fourier transformation (FFT). The results are shown in Figure 2B, 2D and 2F. A peak of power in the theta-band is most noticeable. To quantify the significance of the observed spectral power, we next performed a randomization procedure by shuffling time courses of the MVPA classification accuracy for congruent condition and incongruent condition independently for each participant 1000 times. After each randomization, FFT was conducted on surrogate signals, generating a distribution of spectral power for each frequency point from which we obtained statistical thresholds for evaluating significance. The theta-band oscillations of the priming effects were significant in the FFA, PPA and lMC (*p* < 0.05, multiple comparisons corrected). Results of trials when the probe was a house are shown in Figure 3, no such significant theta-band oscillations were found in the FFA, PPA and lMC (*p* < 0.05, multiple comparisons corrected), consistent with our behavior results in Figure 1B and 1C.

**Figure 3.**
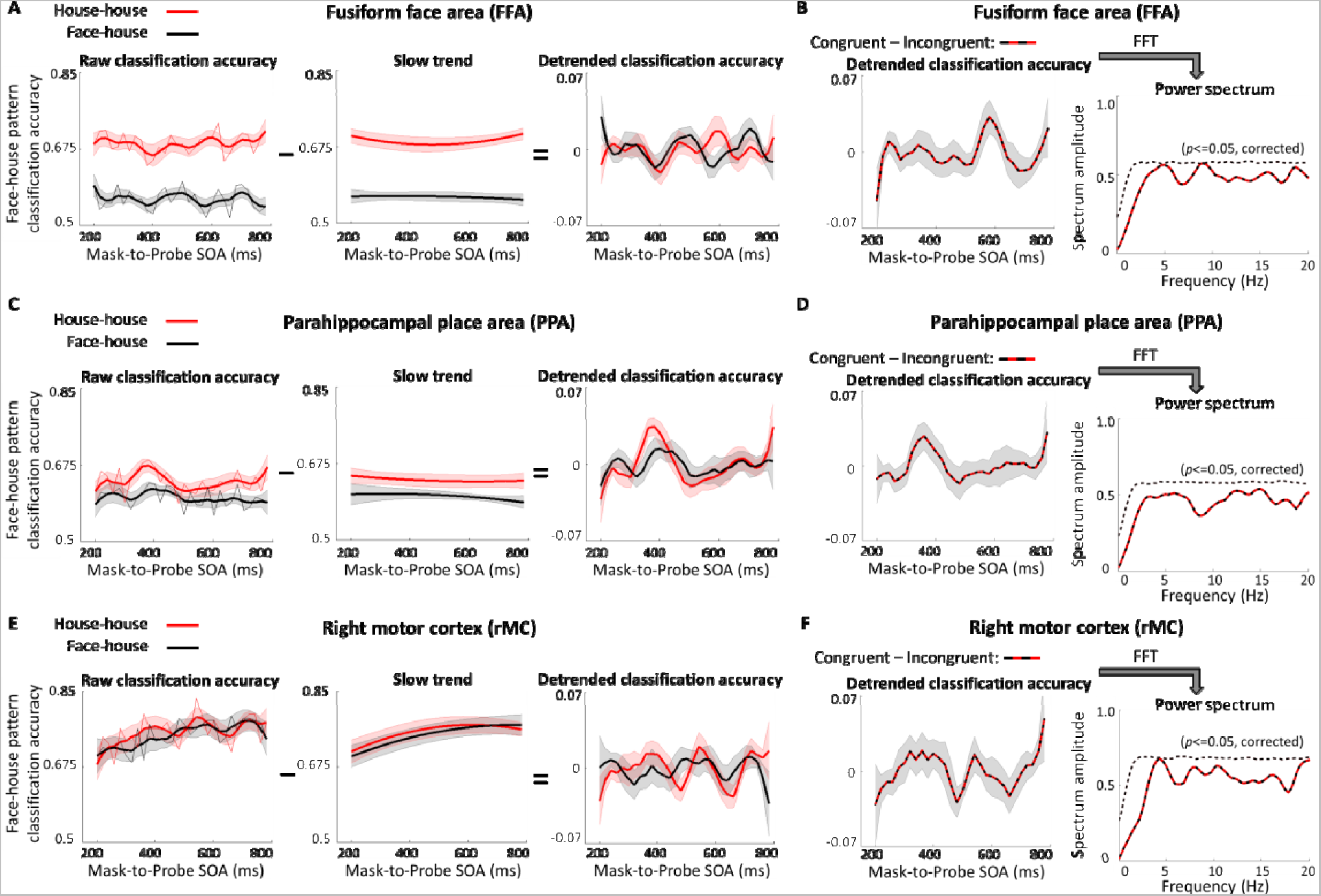
Results of fMRI patterns in the FFA, PPA and lMC when the probe was a house. (A) Results in the FFA. **Left:** Average raw classification accuracies as a function of mask-to-probe SOA (200-780 ms) for congruent (red, thin line) and incongruent (black, thin line) conditions. Average smoothed (60 ms bin) classification accuracies (*n*=18, mean ± SEM) as a function of mask-to-probe SOA, for congruent (red, thick line) and incongruent (black, thick line) conditions. **Middle:** Average slow trends across all the participants. **Right:** Average smoothed-and-detrended classification accuracies extracted by subtracting slow trends shown in ***Middle*** from smoothed (60 ms bin) classification accuracies shown in ***left*** (thick lines). (B) Results of the priming effect (congruent − incongruent) in the FFA. **Left:** Average smoothed-and-detrended classification results of the priming effect. **Right:** Average spectrum for detrended classification accuracies (extracted by subtracting slow trends from the raw classification accuracies) as a function of frequency from 0 to 20 Hz for the priming effect. The statistical threshold of significance (*p* < 0.05, multiple comparison corrected) calculated by performing a permutation test was shown with dashed line. (C) Results in the PPA. (D) Results of the priming effect in the PPA. (E) Results in the rMC. (F) Results of the priming effect in the rMC.

For comparison, no significant theta-band oscillations were found in the rMC, BA17 and ACC (Figure 4). Note that participants used the right hand to report the face probe and the left hand to report the house probe. Given that hand movements are mainly controlled by the contralateral hemisphere, our results for the left and right motor cortices are thus also consistent with the behavioral RT results. Corresponding to that the oscillation in RT time courses can be clearly seen when the probe was a face (Figure 1B), there were significant theta-band oscillations in the lMC (Figure 2F); by contrast, corresponding to no oscillation in RT time courses when the probe was a house (Figure 1C), there were no significant theta-band oscillations in the rMC (Figure 4B). This contrast suggests that our results were unlikely caused some artifacts or data preprocessing.

**Figure 4.**
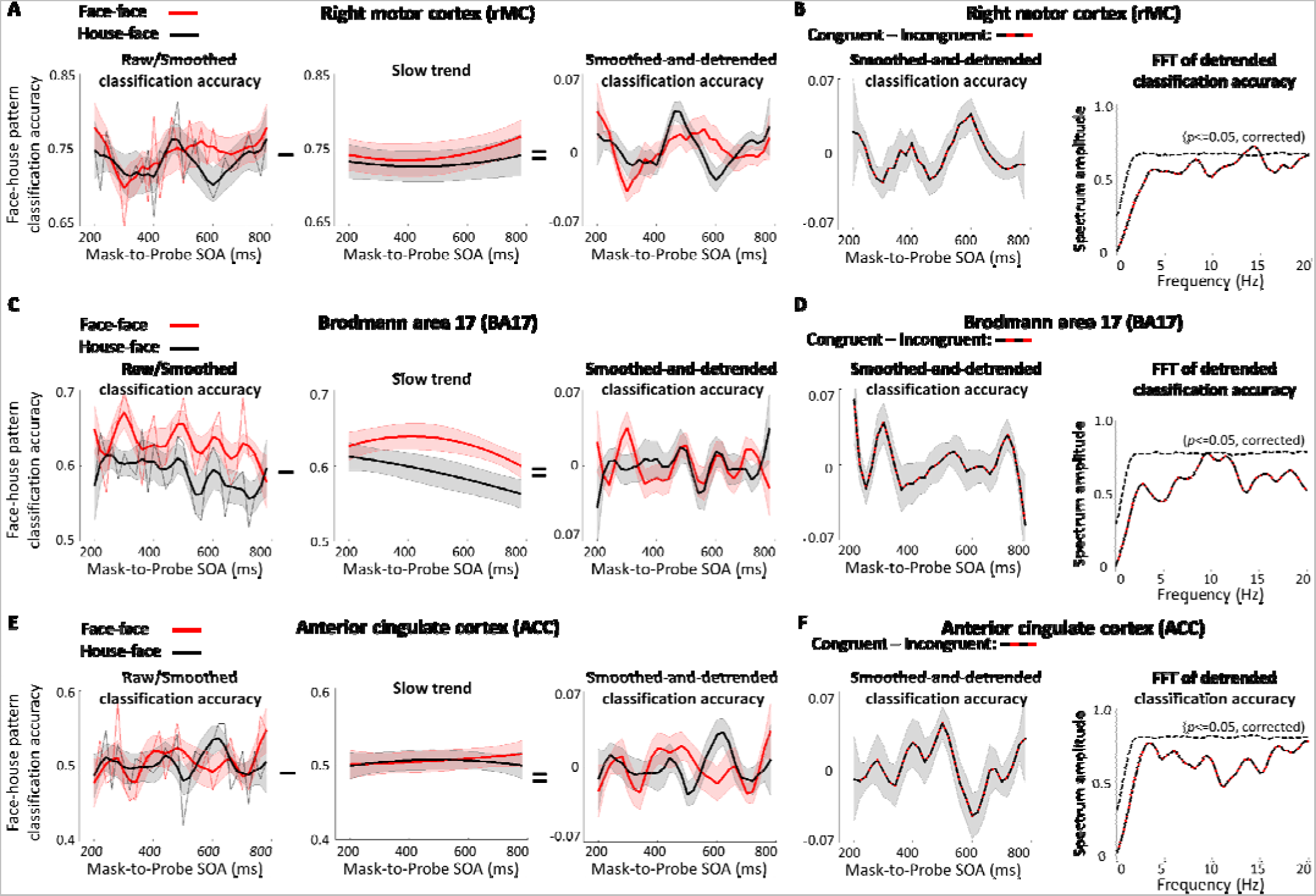
No significant theta-band oscillations in the rMC, BA17 and ACC. (A) Classification results when the probe was a face in the rMC. **Left:** Average raw classification accuracies as a function of mask-to-probe SOA (200-780 ms in steps of 20 ms) for congruent (red, thin line) and incongruent (black, thin line) conditions. Average smoothed (60 ms bin) classification accuracies as a function of mask-to-probe SOA, for congruent (red, thick line) and incongruent (black, thick line) conditions. **Middle:** Average slow trends across all the participants. **Right:** Average smoothed-and-detrended classification accuracies extracted by subtracting slow trends shown in ***Middle*** from smoothed (60 ms bin) classification accuracies shown in ***left*** (thick lines). (B) Classification results of the priming effect (congruent − incongruent) when the probe was a face in the rMC. **Left:** Average smoothed-and-detrended classification results of the priming effect (congruent − incongruent). **Right:** Average spectrum for detrended classification accuracies (extracted by subtracting slow trends from the raw classification accuracies) as a function of frequency from 0 to 20 Hz for the priming effect (congruent − incongruent). The statistical threshold of significance (p < 0.05, multiple comparison corrected) calculated by performing a permutation test was shown with dashed line. (C) Results in the BA17. (D) Results of the priming effect (congruent − incongruent) in the BA17. (E) Results in the ACC. (F) Results of the priming effect (congruent − incongruent) in the ACC.

Nonetheless, to further demonstrate that the oscillatory components in the present study were not introduced by any non-oscillatory artifacts or data preprocessing, we generated 18 sets of non-oscillatory (peak at 400 ms) surrogate data, and performed the identical analysis as how the real data were analyzed. No significant theta-band oscillations were found with the surrogate data (Figure 5).

**Figure 5.**
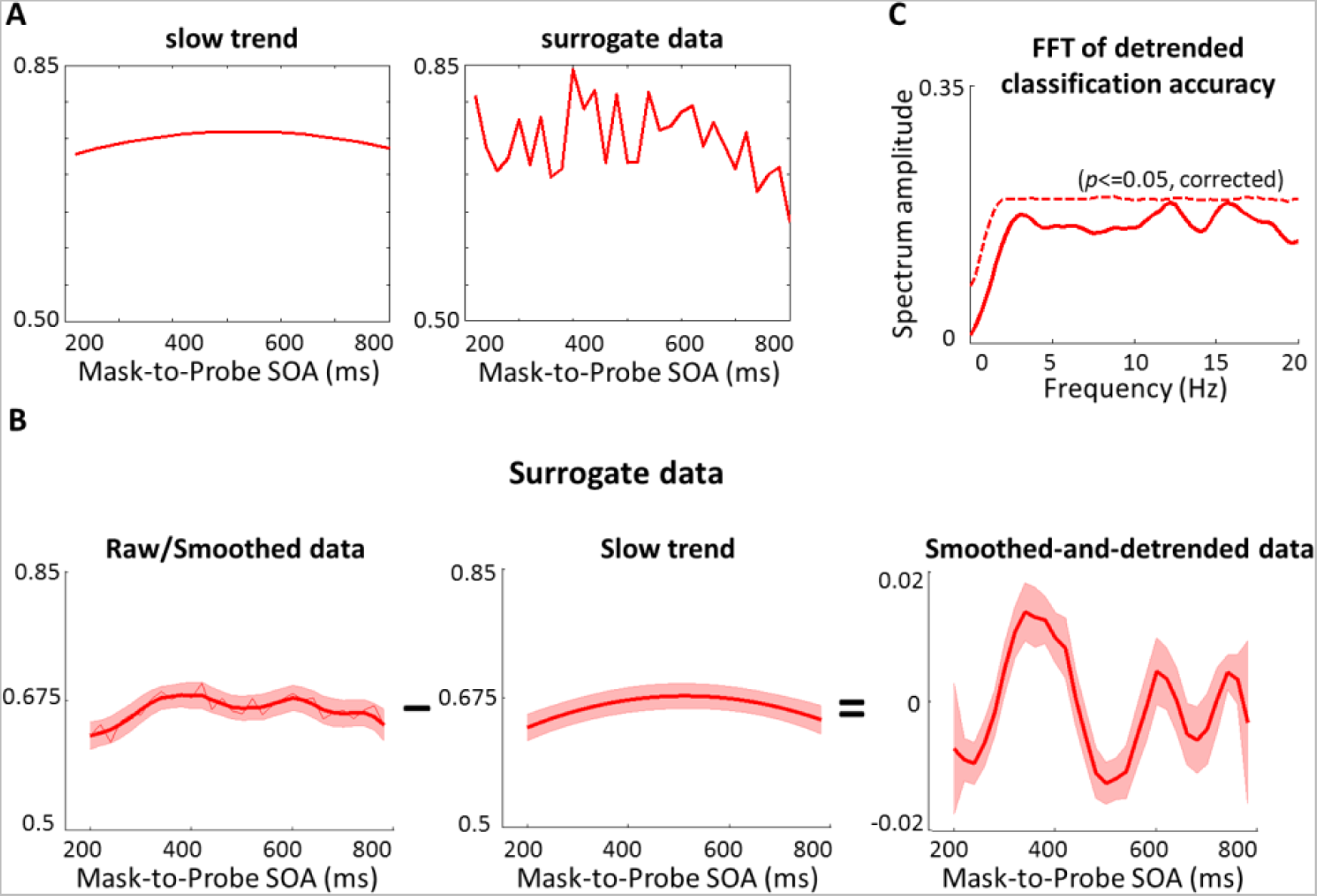
Results of surrogate data. (A) Slow trend and surrogate data of one participant. The slow trend is the slow trend of congruent condition in the FFA, there were 18 different slow trends; the surrogate data were generated by adding a Gaussian curve peaked at 400 ms and white noise (different for each participant) to the slow trend for each participant. Thus, 18 sets of surrogate data were generated. Subsequent analyses of these surrogate data are identical to how we analyzed the real data. (B) **Left:** Averaged surrogate data (*n*=18, mean ± SEM), smoothed (60 ms bin) as a function of mask-to-probe SOA (200-780 ms in steps of 20 ms). **Middle:** Slow trends averaged across participants. **Right:** Average smoothed-and-detrended data, extracted by subtracting slow trends shown in ***Middle*** from smoothed (60 ms bin) data shown in ***left*** (thick lines). (C) Average spectrum for detrended data (extracted by subtracting slow trends from the surrogate data without smoothing). The statistical threshold of significance (*p* < 0.05, multiple comparison corrected) calculated by performing a permutation test was shown with a dashed line.

More interestingly, the theta amplitude in the FFA significantly correlates with the behavioral RT results for the face probe trials across participants, suggesting an important functional role of the theta oscillation in mediating behavior. Specifically, a peak was identified for each participant within the theta-band (3-8 Hz) of the priming effect (congruent – incongruent) oscillations. The amplitude of this peak negatively correlates with the RT in the congruent face-face condition (Pearson’s correlation r = −0.59, *p* < 0.05, Bonferroni corrected) as well as in the incongruent house-face condition (Pearson’s correlation r = −0.62, *p* < 0.05, Bonferroni corrected). Scatter plots of theta oscillation amplitude and RT are shown in Figure 6. Despite moderate subject numbers, a clear trend is visible for greater theta oscillation amplitude in the FFA corresponding to faster reaction time to detect a face probe, suggesting that theta oscillations in the FFA activity patterns may facilitate predictive coding of faces.

**Figure 6.**
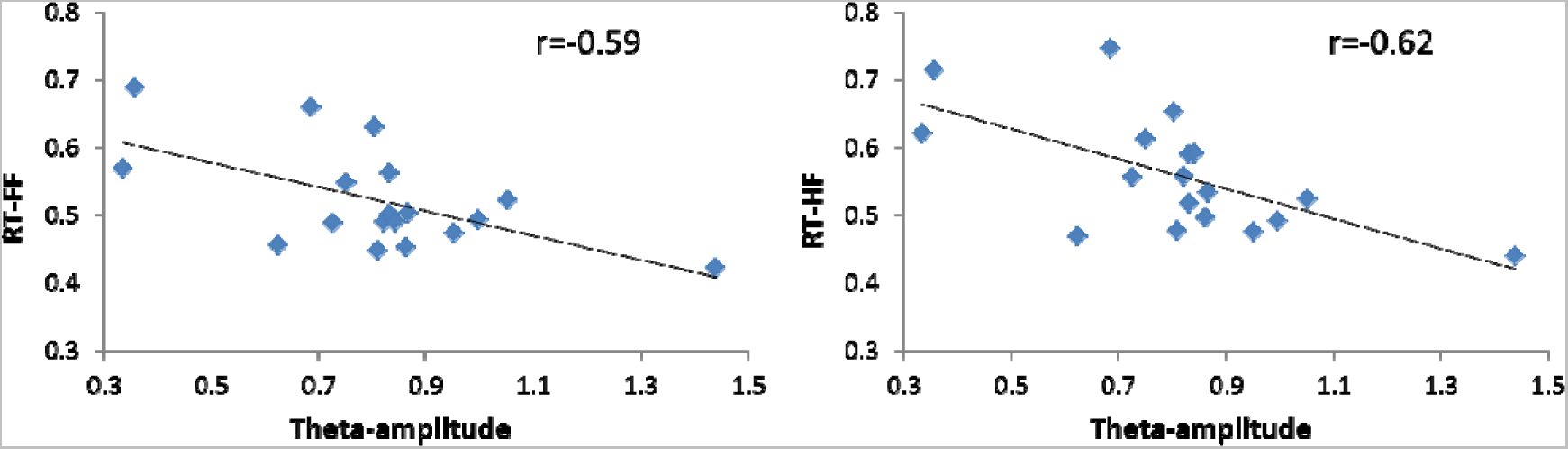
Scatter plots of theta amplitude and RT. The theta amplitude in the FFA negatively correlates with the RT in the FF (face-face) condition (Left: Pearson’s r = −0.59, p < 0.05, Bonferroni corrected) as well as in the HF (house-face) condition (Right: Pearson’s r = −0.62, p < 0.05, Bonferroni corrected).

Results in the left and right motor cortices of a spectrotemporal analysis on detrended MVPA classification accuracies are shown in Figure 7A. This analysis revealed fine dynamic structures of priming effects (congruent − incongruent), as a function of frequency (0-25 Hz) and time (mask-to-probe SOA: 200-780 ms). Significant (permutation test, *p* < 0.05, multiple-comparison corrected across frequencies) theta-band oscillations (~5 Hz) were found in the lMC. By contrast, no significant theta-band oscillations were found in the rMC. Figure 7B, left, shows respectively for congruent (red) and incongruent (black) conditions of the power spectrum of detrended MVPA results in the lMC. Significant theta-band power was found for both the congruent and the incongruent conditions (permutation test, *p* < 0.05, corrected). Interestingly, further phase analysis revealed that the theta-band power of the congruent condition was reliably out of phase with the incongruent condition (Rayleigh test, *p* = 0.03), and clustered around a mean of −160.6° (Figure 7B, right), suggesting a competition-like relationship between the two predictions (face and house) in motor cortex.

**Figure 7.**
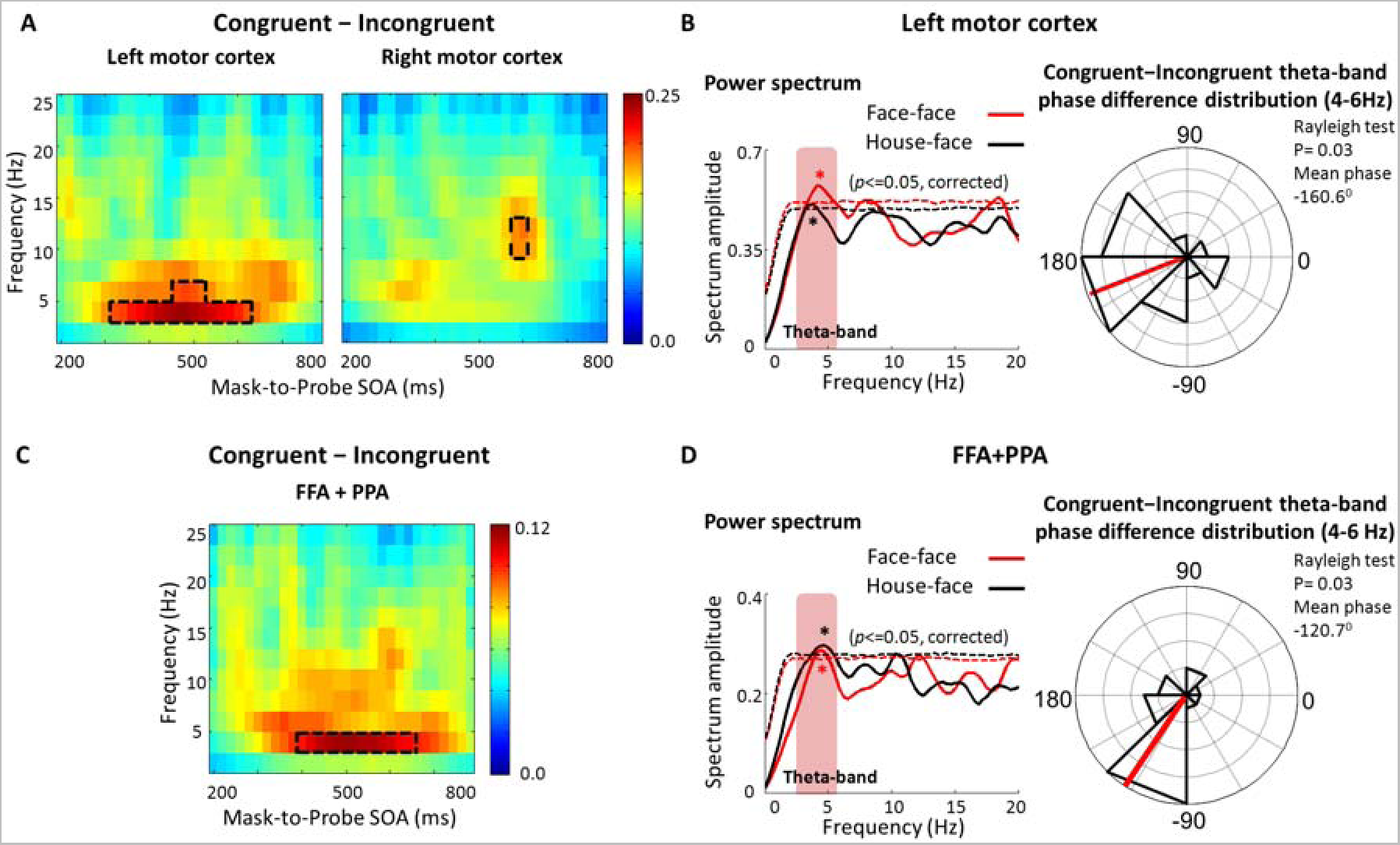
Oscillations of the priming effect (congruent – incongruent). (A) Time-frequency power profiles (*n*=17) for detrended classification accuracies (extracted by subtracting slow trends from the raw classification accuracies) as a function of mask-to-probe SOA (200-780 ms) and frequency (0-25 Hz) in the lMC (left) and rMC (right). Areas enclosed by dashed lines represent statistical significance (*p* < 0.05, multiple comparison corrected) calculated by performing permutation tests. (B) **Left:** Average spectrum for detrended classification accuracies as a function of frequency from 0 to 20 Hz for congruent (red) and incongruent (black) conditions in the lMC. Dashed lines represent the statistical thresholds of significance (*p* < 0.05, multiple comparison corrected) calculated by performing permutation tests. **Right:** Polar plots for the distribution of phase differences between congruent and incongruent conditions in the theta-band (4-6Hz) in the lMC. The red line indicates the mean congruent – incongruent theta-band phase difference across participants. (C) Time-frequency power profile for detrended classification accuracies in the FFA+PPA results. (D) **Left:** Average spectrum for detrended classification accuracies for congruent (red) and incongruent (black) conditions in the FFA+PPA results. **Right:** Polar plots for the distribution of phase differences between congruent and incongruent conditions in the theta-band (4-6Hz).

Moreover, when the FFA and PPA MVPA classification accuracies were combined (Figure 7C and 7D), the theta-band power of the congruent condition was reliably out of phase with the incongruent condition (Rayleigh test, *p* = 0.03), and clustered around a mean of −120.7°. On the one hand, it is possible that combining the FFA and PPA simply increased the statistical power to reveal the out of phase relationship between the congruent and incongruent conditions. On the other hand, this result may imply that decoding face/non-face probe detections could have been benefited from MVPA of not only the FFA but also the PPA -- as for example, while patterns in the FFA may encode faces, patterns in the PPA may contribute to decode that a house (thus not a face) was detected.

### Distinct cortical distributions of theta and alpha oscillations

To further examine the cortical distribution of theta-band oscillations, a whole-brain searchlight analysis was conducted. For each participant, voxels were extracted from a spherical searchlight with a two-voxel radius (33 voxels in each searchlight including the central voxel), and then MVPA was performed using this spherical searchlight ROI, which moved throughout each participant’s whole brain gray matter-masked data. Frequency analyses were conducted for each searchlight ROI to calculate the power of theta-band oscillations, and then the results were assigned to the central voxel of the sphere searchlight. After normalization (Z-score) across all voxels, clusters with significant power (*p* < 10^−4^) and size (>15 voxels) were localized. Thus, a significant cluster indicates that MVPA classification accuracy of the probe fluctuated as a function of SOA in the theta band.

For comparisons, the same searchlight procedure was also performed to map clusters (>15 voxels, except one subject >11 voxels, *p* < 10^−4^) of significant alpha-band (8-13 Hz) oscillations across the whole brain. Abundant EEG and MEG studies have demonstrated that alpha-band oscillations were predominantly observable in occipital sites (Thut et al. 2006), and we are the first to map the alpha-band oscillations using fMRI. Alpha oscillations correlate with cortical inhibition (Ray and Cole 1985, Palva and Palva 2007), and the ongoing occipital alpha oscillations have been argued to play direct functional roles in attention and perception mechanisms (VanRullen and Koch 2003, Busch and VanRullen 2010, Dugue, Marque, and VanRullen 2011, Jensen, Bonnefond, and VanRullen 2012). Moreover, the cross-frequency coupling between alpha and theta oscillations has been reported recently by using time-resolved RT measurements (Huang, Chen, and Luo 2015, Song et al. 2014).

Figure 8A shows percentages (Y-axis) of participants, in whom significant clusters were found based on the searchlight analysis in the occipital, temporal, parietal, and frontal lobes. Remarkably, at least one cluster of theta oscillations was reliably found in the temporal cortex in all participants (100%), whereas at least one cluster of alpha oscillations was reliably found in the occipital cortex of all participants (100%). By contrast, this level of concentration was not seen for parietal and frontal cortices, given the criterion we used to localize the clusters is fairly stringent and on average only 2.5 clusters for theta oscillations and 4.3 clusters for alpha oscillations were found per each participant. Using Freesurfer and an atlas-based automatic surface parcellation (Desikan et al. 2006, Fischl et al. 1999, Fischl et al. 2004), spatial distributions of brain regions that exhibited significant theta-band and alpha-band power are further shown in Figure 8B and 8C, respectively. The magnitude of the color scale in Figure 8B and 8C indicates the number of significant clusters per 1000 mm^2^ that were found in the marked atlas-based anatomical regions of interest. Gray-colored areas indicate there was no significant cluster, whereas yellow indicates there were ~4 significant clusters per 1000 mm^2^ in each of the marked parcellated cortical regions. Differences between the two distributions are obvious: most of the regions with significant theta-band oscillations were in the temporal cortex, and most of the regions with significant alpha-band oscillations were in the occipital cortex.

**Figure 8.**
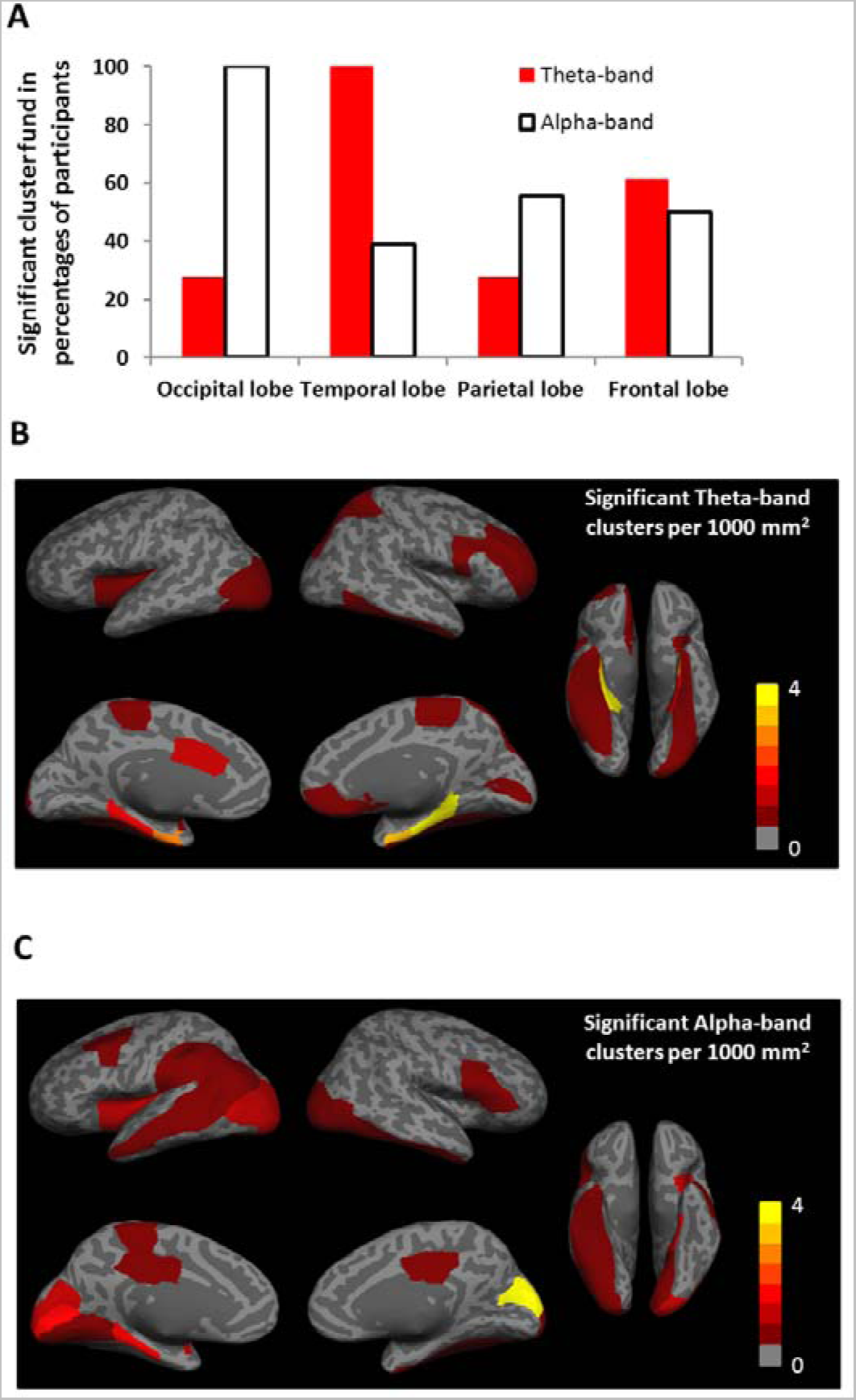
Different spatial distributions of theta-band and alpha-band oscillations. (A) Y-axis indicates the percentages of participants, in whom significant clusters were found based on the searchlight analysis in the occipital, temporal, parietal, and frontal lobes. Significant clusters of theta-band oscillations were found in the temporal lobe of every participant (100%), whereas significant clusters of alpha-band oscillations were found in the occipital lobe of every participant (100%), far more concentrated and robust than any other lobes. (B) The cortical distribution map of theta-band oscillations. The magnitude of the color scale indicates the number of significant clusters per 1000 mm^2^ that were found in the marked atlas-based anatomical regions of interest. Gray-colored areas indicate there was no significant cluster, whereas yellow indicates there were ~4 significant clusters per 1000 mm^2^ in each of the marked parcellated cortical regions. (C) The cortical distribution map of alpha-band oscillations.

## DISCUSSION

These combined psychophysical and neuroimaging results provide important constraints on the hypothesized links between theta rhythms and dynamic predictive coding. Multi-voxel activation patterns in the FFA and PPA have been suggested to encode object categories (Haxby 2012, Haxby et al. 2001, Haynes and Rees 2006), and here we also demonstrated classical priming effects in FFA and PPA. More interestingly, however, our study revealed that multi-voxel activation patterns in these brain areas are not stationary but fluctuate as a function of mask-to-probe relation at a theta-band rhythm. This fluctuation is sustained across the tested SOA periods over which we can discount hemodynamic lags, since we were comparing trial-by-trial differences as a function of mask-to-probe SOA, assuming that hemodynamic lags in a same ROI are always comparable across trials. Thus, rather than bottom-up perceptual responses to the incoming stimulus, our findings more likely reveal effects of the memory traces (priming) that mediate predictive coding. This result is consistent with the hypothesis that perception is modulated by ongoing theta oscillations whose phase is reset by priming (Huang, Chen, and Luo 2015, Busch, Dubois, and VanRullen 2009, Song et al. 2014, Romei, Gross, and Thut 2012), but, crucially, is the first to show that theta rhythms in the fluctuation of multi-voxel activity patterns are linked to predictive coding effects. Indeed, greater theta oscillation amplitude in the FFA significantly correlated to faster reaction time to detect a face probe, directly supporting the functional link.

By using fMRI and a whole-brain searchlight analysis, we were further able to map more precisely the cortical distributions of various brain rhythms. Alpha oscillations were concentrated in the occipital lobe, which is consistent with previous EEG and MEG reports (Thut et al. 2006), whereas theta oscillations were concentrated in the temporal lobe, suggesting distinct functional roles theta-band oscillations may play. Moreover, given that an fMRI voxel may contain millions of neurons (Logothetis 2008), fluctuations of activity of a small number of neurons are unlikely to cause fluctuations of multi-voxel fMRI activity patterns. What we have observed through fMRI and MVPA thus presumably reflects rhythmic ensemble responses across distributed populations of neurons (Haxby 2012, Kriegeskorte et al. 2008, Guo and Meng 2015).

Note that the temporal profiles of FFA, PPA and lMC are different in phase and frequency, suggesting that the observed theta oscillations were unlikely underlying possible long-range coordination of activity in these brain areas. We thus propose that priming leads to a reset of ongoing theta-band oscillations, which were recently reported to be involved in attention and predictive coding (Huang, Chen, and Luo 2015, Landau and Fries 2012, Fiebelkorn, Saalmann, and Kastner 2013, Song et al. 2014). And because the SOA varied in small steps (20 ms), the subsequent probe was processed at different phase of this reset oscillation, enabling us to observe the periodic neuroimaging pattern. To rule out the possibility that artifacts or data preprocessing could have introduced oscillatory signatures into our results, we generated sets of surrogate non-oscillatory data, and performed the exact analysis procedure as how the real data was analyzed. No significant theta-band oscillations were found with the surrogate data (Figure 5). It remains possible that the fine-scale temporal profiles we found in the FFA, PPA and lMC were not really oscillatory, but only peaking a few times within the limited SOA range that we had examined. Future studies can further investigate this possibility by using a longer SOA range. Nevertheless, the present study revealed relatively fine-scale temporal dynamics of fMRI activity patterns.

Our study provides a feasible strategy that incorporates fMRI and MVPA to investigate the dynamics of ensemble coding across distributed populations of neurons. Despite the hemodynamic lag, fMRI has been combined with novel behavioral paradigms to probe neural responses at the time scale of tens of milliseconds, since neuronal electrical activity and the fMRI signal are reliably coupled at this level of temporal precision (Ogawa et al. 2000, Formisano et al. 2002, Dux et al. 2006). Specifically, one of these previous studies (Dux et al. 2006) compared fMRI responses in two different SOA conditions (a short SOA and a long SOA), demonstrating that fMRI can be used to measure temporal dynamics of visual processing.

MVPA has been shown to have better temporal resolution than the univariate measurement of BOLD activity change (Kohler et al. 2013). It is therefore expected that the time-resolved strategy we advocate should be more sensitive than conventional fMRI approaches to detect the dynamics of trial-by-trial fluctuation in population coding as a function of SOA. Consistent with this notion, we conducted the same spectrum analysis with univariate averaged BOLD responses in the FFA, PPA, and lMC (Figure 9). There were no significant theta-band oscillations in the univariate averaged FFA or PPA activity. And while theta-band oscillation was found in the lMC, it did not show significant out-of-phase relationship between the congruent and incongruent conditions (Rayleigh test, *p* = 0.17). These results differ from the MVPA results shown in Figure 2 and Figure 7, confirming that multi-voxel fMRI activity patterns instead of merely averaged fMRI activity fluctuate.

**Figure 9.**
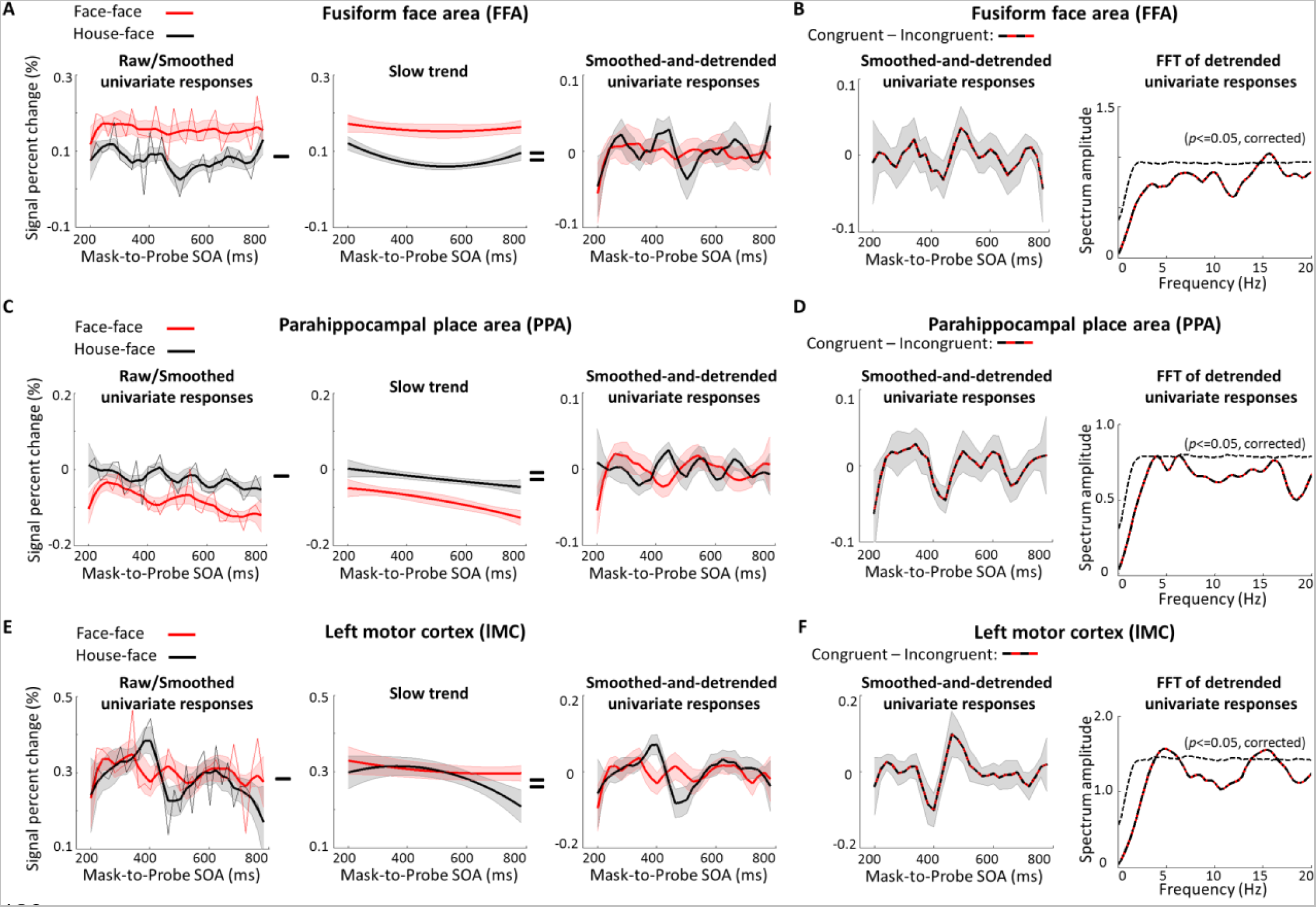
Results of univariate averaged BOLD responses in the FFA, PPA and lMC. (A) Results of when the probe was a face in the FFA. **Left:** Averaged BOLD responses as a function of mask-to-probe SOA (200-780 ms in steps of 20 ms) for congruent (red, thin line) and incongruent (black, thin line) conditions. Smoothed (60 ms bin) BOLD responses (*n*=18, mean ± SEM) as a function of mask-to-probe SOA, for congruent (red, thick line) and incongruent (black, thick line) conditions, show an overall priming effect (the congruent condition evoked overall greater BOLD responses than the incongruent condition). **Middle:** Average slow trends across all the participants. **Right:** Average smoothed-and-detrended BOLD responses extracted by subtracting slow trends shown in ***Middle*** from smoothed (60 ms bin) BOLD responses shown in ***left*** (thick lines). (B) Results of the priming effect (congruent − incongruent) when the probe was a face in the FFA. **Left:** Average smoothed-and-detrended BOLD results of the priming effect (congruent − incongruent). **Right:** Average spectrum for detrended BOLD responses (extracted by subtracting slow trends from the raw BOLD responses) as a function of frequency from 0 to 20 Hz for the priming effect (congruent − incongruent). The statistical threshold of significance (*p* < 0.05, multiple comparison corrected) calculated by performing a permutation test was shown with dashed line. (C) Results in the PPA. (D) Results of the priming effect (congruent − incongruent) in the PPA. (E) Results in the lMC. (F) Results of the priming effect (congruent − incongruent) in the lMC. While a significant 15-17 Hz component can also be generally seen in B, D, and F, as well as B in S1 Figure and B in Figure 2, it may be caused by artifacts from acoustic noises generated by our EPI sequence (17.5 Hz). We thus choose to not discuss this 15-17 Hz component further in the main text.

To conclude, we combined fMRI and a time-resolved psychophysical paradigm to investigate the dynamic neural mechanism underlying visual object priming. Specifically, multi-voxel activity patterns in the FFA and the PPA show temporal fluctuations at a theta-band (~5 Hz) rhythm, suggesting the critical role of theta oscillations in the inferior temporal cortex during visual object priming. Our strategy is obviously not limited to the theta-band and predictive coding, and future studies may take similar approaches to better understand other mechanisms underlying brain dynamics.

### Materials and methods

#### Participants

Eighteen healthy adults (7 females; mean age 26 years; all right handed) participated in this two-session fMRI experiment. All participants had normal or corrected-to-normal visual acuity and gave written informed consent. This study was approved by the Dartmouth College Committee for the Protection of Human Subjects.

#### MRI acquisition

Participants were scanned using a 3T Philips Achieva Intera scanner with a 32-channel head coil at the Dartmouth Brain Imaging Center. An echo-planar imaging (EPI) sequence (2000 ms TR; 35 ms TE; 3 × 3 × 3 mm voxel size; 35 slices) was used to measure the BOLD contrast. For each participant, a high-resolution T1-weighted anatomical scan was acquired at the beginning (or the end) of each scan session (8.2 ms TR; 3.8 ms TE; 1 × 1 × 1 mm voxel size; 222 slices). During the EPI scans, visual stimuli were presented to a screen located at the back of the scanner via a LCD projector (Panasonic PT-D4000U) using MATLAB 2011b with Psychtoolbox(Brainard 1997). Participants viewed the stimuli using a mirror placed within the head coil.

#### Region of Interest (ROI) localizer runs

An independent set of gray-scale face and house images was used to localize the ROIs. The localizer scans consisted of an alternating block design, with 5 blocks presenting face images and 5 blocks presenting house images interleaved with 16-s periods of a blank screen with a fixation cross in the center of the screen. Each stimulus block was also 16-s long. In total, each localizer scan run was 336-s long, consisting of 11 periods of fixation and 10 stimulus blocks. In each stimulus block, 16 faces (or houses) were presented (500 ms per image, with a 500-ms interstimulus interval). Fifteen participants completed two localizer scans, and three participants completed three localizer scans. During localizer scans, participants performed a new face (or house) detection task in which they were asked to use their right hand to make a key-press whenever a new face was presented and their left hand to make a key-press whenever a new house was presented. Twelve new faces/houses were presented in each block. This task also allowed us to localize the left and right motor cortices as ROIs by contrasting BOLD responses corresponding to right-hand button presses vs. left-hand button presses.

#### Experimental runs

Each participant completed 24 experimental scan runs in two sessions on two separate days. Each experimental scan run was 368-s long, consisting of 90 trials and two 4-s periods (at the beginning and the end of each run) of a blank screen with a fixation cross in the center of the screen. As shown in Figure 1A, each trial was 4-s long, and presented in a semi-randomized order with a rapid event-related design. In each trial, a 150-ms probe was preceded by a 33-ms prime stimulus which was backward masked by a 100-ms mask stimulus. Critically, the mask-to-probe SOAs ranged from 200 to 780 ms in steps of 20 ms, corresponding to a sampling frequency of 50 Hz. The stimuli included two images, a high-contrast face and a low-contrast house. The high contrast level was defined with root mean square (RMS) = 0.25 in normalized unit, whereas the low contrast level RMS = 0.025. Corresponding contrasts of the face image and the house image were made by using the SHINE toolbox (Willenbockel et al. 2010). The probe and the prime were either the same (congruent conditions: Face-prime followed by Face-probe; House-prime followed by House-probe) or different (incongruent conditions: Face-prime followed by House-probe; House-prime followed by Face-probe), except that the probe (5.8°) was smaller than the prime (8.7°) to avoid any possible low-level effects of retinotopic adaptation (Figure 1A). In total, there were 2160 trials for each participant: 12 repetitions for each of the four prime-probe conditions at each of the 30 SOAs (from 200 to 780 ms in steps of 20 ms). Participants were asked to report whether the probe was a face (right hand button press) or a house (left hand button press) with speeded responses.

#### MR Image preprocessing

AFNI (Cox 1996)(<u>http://afni.nimh.nih.gov/afni)</u> was used for preprocessing the MRI data. EPIs were slice timing corrected, motion corrected to the image acquired closest to the anatomical images, spatially smoothed with a 4-mm full width at half maximum filter (FWHM), and temporally filtered to remove baseline drifts. Based on the anatomical images acquired in each of the two sessions, mean anatomical images were computed to remove the bias of either session. All EPIs were then aligned to the mean anatomical images.

#### Functional ROI localization

Data from the ROIs localizer scans were further submitted to a General Linear Model (GLM) analysis, which calculated the beta coefficient values associated with block conditions. ROIs were individually defined for each participant based on activation maps from the GLM analysis. Among them, four were defined as a continuous cluster of activated voxels corresponding to the following GLM contrasts: the FFA was defined in the middle fusiform gyrus as responding more strongly to faces than to houses; the PPA was defined in the parahippocampal gyrus as responding more strongly to houses than to faces; the left motor cortex was defined as responding more strongly to right-hand button presses than to left-hand button presses; and the right motor cortex was defined as responding more strongly to left-hand button presses than to right-hand button presses. To control for any potential confounding effects of ROI size, the statistical contrast threshold was adjusted individually (maximum *p* < 10^−4^, uncorrected) to roughly match the size of each of these ROIs (~40 voxels). Using this threshold, however, did not allow for the localization of the motor cortex ROI in one participant. Therefore, subsequent ROI analysis of the motor cortex did not include this participant. Next, data from the ROI localizer scan runs were aligned to Talairach space using the TT_N27 template. For each participant, Brodmann area 17 (BA17) was localized using an anatomical mask based on TT_N27 template as well as GLM activation maps, which include activated voxels in the calcarine sulcus that responded more strongly during stimulation blocks than during fixation periods (maximum *p* < 10^−4^, uncorrected). The size of the BA17 ROI was on average 100 voxels. Similarly, data from the experimental scan runs were aligned to the Talairach space (TT_N27 template) and were submitted to a GLM analysis to calculate the beta coefficient values associated with congruent conditions (Face-prime followed by Face-probe; House-prime followed by House-probe) and incongruent conditions (Face-prime followed by House-probe; House-prime followed by Face-probe). The GLM activation map corresponding to congruent vs. incongruent differences (*p* < 10^−2^, uncorrected, ~40 voxels) was used to localize the ROI of anterior cingulate cortex (ACC).

#### Univariate averaged BOLD response and multivariate pattern classification analyses

For each participant, we extracted the averaged activation values across all voxels in each ROI to analyze the univariate BOLD response changes. Percent BOLD signal change was calculated for each trial by using the average of the last TR before and the first TR after the trial onset as the baseline. Subsequent analyses focused on the peak percent signal change amplitude at the third TR (6s) from the trial onset. Multivariate pattern analyses (MVPA) were performed using PyMVPA (Hanke et al. 2009). We extracted the activation values of all voxels in each ROI for each trial, removing the mean intensity of the ROI, to compute the multi-voxel activation pattern based on the third TR (6s) from the trial onset. Pattern classifications of the face probe condition and the house probe condition were then performed with linear support vector machines (SVMs) using a leave-one-trial-out cross-validation procedure.

#### Analyses of frequency and phase

Analyses of frequency and phase were performed with MATLAB (The MathWorks) using functions from the EEGLAB toolbox (Delorme and Makeig 2004) and CircStat toolbox (Berens 2009). First, we calculated the temporal profile of RTs/averaged BOLD responses/MVPA classification accuracies as a function of SOA from 200 to 780 ms in steps of 20 ms (50 Hz sampling frequency) for each condition (congruent and incongruent). For each participant, in order to extract the slow developing trend, raw RTs/averaged BOLD responses/MVPA classification accuracies of each condition were fitted to a second order polynomial function. We then subtracted the slow trend from corresponding temporal profile for each participant to obtain detrended RTs/averaged BOLD responses/MVPA classification accuracies separately for each condition to remove possible interferences from classical priming and expectancy effects. Next, to further investigate the oscillatory patterns of priming effects, we subtracted the detrended temporal profiles of incongruent conditions from congruent conditions. To investigate the spectral characteristics of the detrended priming effects, we then conducted spectrum analysis separately for each participant. Specifically, we performed a Fast Fourier transformation (FFT) to convert the detrended priming effects into the frequency domain (after zero padding and application of a Hanning window). In this study the FFT length was 160 data points and the window size was 40 data points. Also, to examine the phase relationships between the congruent and incongruent conditions, testing for nonuniformity for congruent − incongruent phase differences in the theta-band (4-6 Hz) across participants was conducted using circular statistics (Rayleigh test for nonuniformity for circular data in CircStats toolbox). We further performed a randomization procedure by shuffling the RTs/averaged BOLD responses/MVPA classification accuracies for congruent condition and incongruent condition respectively within each participant to assess the statistical significance of the observed spectral power as well as the congruent − incongruent phase relationship. After each randomization, we conducted FFT on surrogate signals, similar to that of the original data analysis; we repeated this procedure 1000 times, arriving at a distribution of spectral power for each frequency point from which we obtained the *p* < 0.05 threshold (uncorrected). We then applied multiple comparison correction to the uncorrected randomization threshold spectrum profile. Similarly, for each randomization, we conducted the same phase analysis on the surrogate signals by calculating cross-participant coherence in the congruent − incongruent phase difference.

#### Time-frequency analysis

To assess MVPA classification accuracies as a function of time (mask-to-probe SOA) and frequency, the detrended temporal profile for each condition was transformed using the continuous complex Gaussian wavelet (order = 4; e.g., FWHM =1.32 s for 1 Hz wavelet) transforms (Wavelet toolbox, MATLAB), with frequencies ranging from 1 to 25 Hz in steps of 2 Hz. The power profile of detrended classification accuracies (squared absolute value) as a function of time and frequency was then extracted from the output of the wavelet transform. Power profiles for priming effects (congruent − incongruent) were calculated for each participant separately. The grand mean of time-frequency power was then calculated by averaging across participants. We further performed a randomization procedure to assess the statistical significance of the power profiles for priming effects (congruent − incongruent), by shuffling the labeling of SOAs. After each randomization, the same time-frequency analysis was performed on the surrogate signals, as that performed in the original data analysis. This procedure was repeated 1000 times and resulted in a distribution of power at each time-frequency point, from which the *p* < 0.05 threshold (uncorrected) was obtained. The cross-frequency multiple-comparison correction was then further applied to the uncorrected randomization threshold time-frequency map.

#### Whole-brain searchlight analysis

A whole brain searchlight analysis (Kriegeskorte, Goebel, and Bandettini 2006)was conducted to identify brain regions where significant theta-band (3-6 Hz) and alpha-band (8-11 Hz) oscillations occurred. For each participant, voxels were extracted from a spherical searchlight with a two-voxel radius (33 voxels in each searchlight including the central voxel) and then MVPA was performed using this spherical searchlight ROI. The searchlight moved throughout each participant’s gray matter-masked data using PyMVPA (Hanke et al. 2009). For each searchlight corresponding to a central voxel (i.e., each voxel across the whole gray matter mask), a linear SVM learning algorithm was trained and tested to examine pair-wise classification performance for face probe vs. house probe conditions. To ensure independence between training and testing, cross-validations were performed between even scan runs and odd scan runs (train on even runs, test on odd runs; and vice versa). Next, frequency analyses were conducted for each searchlight ROI to calculate the power of theta-band and alpha-band oscillations, and then the results were assigned to the central voxel of the sphere searchlight. After normalization (Z-score) across all voxels, clusters with significant power (*p* < 10^−4^) and size >15 voxels (except one subject for alpha-band cluster >11 voxels) were localized. Percentages of participants with clusters in the temporal, parietal, frontal and occipital lobes were calculated separately for theta-band and alpha-band oscillations. Further, to show the spatial distribution of brain regions that exhibited significant theta-band and alpha-band power, Freesurfer was used to generate atlas-based automatic surface parcellation and to map the concentration by numbers of theta and alpha oscillation clusters per 1000 mm^2^ in each of the parcellated cortical regions.

## ACKNOWLEDGMENTS

This work is supported by the National Science Foundation under Grant No. 1157121.

